# Activation of Piezo1 but not Na_V_1.2 Channels by Ultrasound at 43 MHz

**DOI:** 10.1101/136994

**Authors:** Martin Loynaz Prieto, Kamyar Firouzi, Butrus T. Khuri-Yakub, Merritt Maduke

**Author notes:** Corresponding Author: Merritt Maduke, Beckman Center B155 279 Campus Drive West Stanford University School of Medicine Stanford, CA 94306.

## Abstract

Ultrasound (US) can modulate the electrical activity of the excitable tissues but the mechanisms underlying this effect are not understood at the molecular level or in terms of the physical modality through which US exerts its effects. Here we report an experimental system that allows for stable patch-clamp recording in the presence of US at 43 MHz, a frequency known to stimulate neural activity. We describe the effects of US on two ion channels proposed to be involved in the response of excitable cells to US: the mechanosensitive Piezo1 channel and the voltage-gated sodium channel Na_V_1.2. Our patch-clamp recordings, together with finite-element simulations of acoustic field parameters indicate that Piezo1 channels are activated by continuous wave US at 43 MHz and 50 or 90 W/cm^2^ through cell membrane stress caused by acoustic streaming. Na_V_1.2 channels were not affected through this mechanism at these intensities, but their kinetics could be accelerated by US-induced heating.

## INTRODUCTION

It has long been known that ultrasound (US) can modulate electrical activity in excitable tissues (Fry, et al. 1958, Gavrilov, et al. 1996, Harvey 1929). In recent years, several research groups have investigated this effect with the motivation of developing US neuromodulation as a tool for treating mental and neurological disorders (Bystritsky, et al. 2011, Kim, et al. 2014, Kim, et al. 2015, King, et al. 2013, King, et al. 2014, Lee, et al. 2015, Lee, et al. 2016, Legon, et al. 2014, Mehic, et al. 2014, Menz, et al. 2013, Min, et al. 2011, Tufail, et al. 2010, Tyler, et al. 2008, Ye, et al. 2016, Yoo, et al. 2011, Younan, et al. 2013). A distinct advantage of US in this context is its ability to be focused deep within tissue with excellent spatial resolution and to function without surgical implants or genetic manipulation. These possibilities motivate the investigation of the mechanistic basis of US effects on excitable tissues.

Despite increased interest in recent years, models and hypotheses for the mechanism of US effects on excitability (Krasovitski, et al. 2011, Plaksin, et al. 2016, Plaksin, et al. 2013, Prieto, et al. 2013, Sassaroli and Vykhodtseva 2016, Tyler 2011) have been much more abundant than definite evidence in support of any particular mechanism. Complicating the picture is the essentially polymodal nature of US interaction with biological tissue (O’Brien 2007). In addition to changes in density and particle displacement related to the primary acoustic pressure, there may also be effects related to acoustic cavitation, temperature changes due to acoustic energy absorption, and second-order effects of radiation force and acoustic streaming. In different clinical and experimental contexts, and for different sets of US stimulus parameters, different modalities may be of primary importance. Clarity could be provided by a system for studying the behavior of the basic “units” of biological electrical activity–individual ion channels–in the presence of US. The gold standard for this type of investigation is patch-clamp recording, which allows for detailed characterization of ion channel kinetics and membrane voltage dynamics. However, published accounts (Tyler, et al. 2008) and our own experience indicate that the gigaOhm seals required for patch-clamp recording are unstable in presence of US in the typical low frequency range used for in vivo neuromodulation (~0.2-3 MHz), and are irreversibly damaged at relatively low US intensities (> 1 W/cm^2^). This instability, however, appears to be frequency dependent. Here we report an experimental system that allows stable patch-clamp recording in the presence of US at 43 MHz, at intensities known to modulate neural activity in the salamander retina *in vitro* (Menz, et al. 2013) and in acute rat hippocampal brain slices *in vitro* (Prieto, et al. 2016).

Using this system, we investigated the effects of US on heterologously expressed mechanosensitive Piezo1 channels and on Na_V_1.2 voltage-gated sodium channels. The choice of these two channels was guided by the hypothesis that US modulates action potential firing *in vivo* by causing membrane stress, thereby affecting the activity of mechanosensitive ion channels. Piezo channels are one of the few eukaryotic channels known to be directly activated (as opposed to modulated) by membrane stress (Coste, et al. 2010, Coste, et al. 2012), and since their discovery in 2010 have come to be regarded as the principal mechanoreceptor channel in mammalian cells (Volkers, et al. 2015, Wu, et al. 2016). In the context of US neuromodulation, it is notable that Piezo channels are expressed in the brain at the messenger RNA level (Coste, et al. 2010). Voltage-gated sodium channels (Na_V_ channels) are a strong candidate for the molecular target of US neuromodulation because of their central role in action potential generation, and because they can be modulated by membrane stress (Beyder, et al. 2010, Morris and Juranka 2007, Wang, et al. 2009).

## MATERIALS AND METHODS

### Experimental set-up

The experimental *set-up* for simultaneous US stimulation and patch-clamp recording of cultured cells is illustrated in Figure 1A. The set-up is based on a modified Axioskop-2 microscope (Zeiss Microscopes, Jena, Germany) with a 40x W N-Achroplan objective (Zeiss Microscopes, Jena, Germany). The condenser was removed and replaced with a custom-built housing for the 43-MHz transducer used in these experiments. US is transmitted from below the cells in the direction perpendicular to the bottom of the experimental chamber. Cells are illuminated with a ring of white LED lights mounted on the transducer housing, allowing visualization of the cells through the epifluoresence pathway with an image of sufficient quality for patch-clamp recording. The transducer is coupled to the bottom of the chamber by a drop of distilled water held by a small rubber O-ring attached to the tip of the transducer with silicone grease. The experimental chamber containing the cells was made from the lid of a 35-mm cell culture dish with a ~1 cm hole in the center. A thin film of plasma-treated polystyrene (Goodfellow USA, Coraopolis, PA, USA) with cultured cells adhering to it was sealed onto the bottom of the chamber using silicone grease. Polystyrene films were used as the cell substrate instead of standard tissue culture dishes in order maximize US transmission and minimize heating due to the absorption of acoustic energy. The thickness of the film was 25 microns, except where indicated in the figure legend.

**Figure 1.**
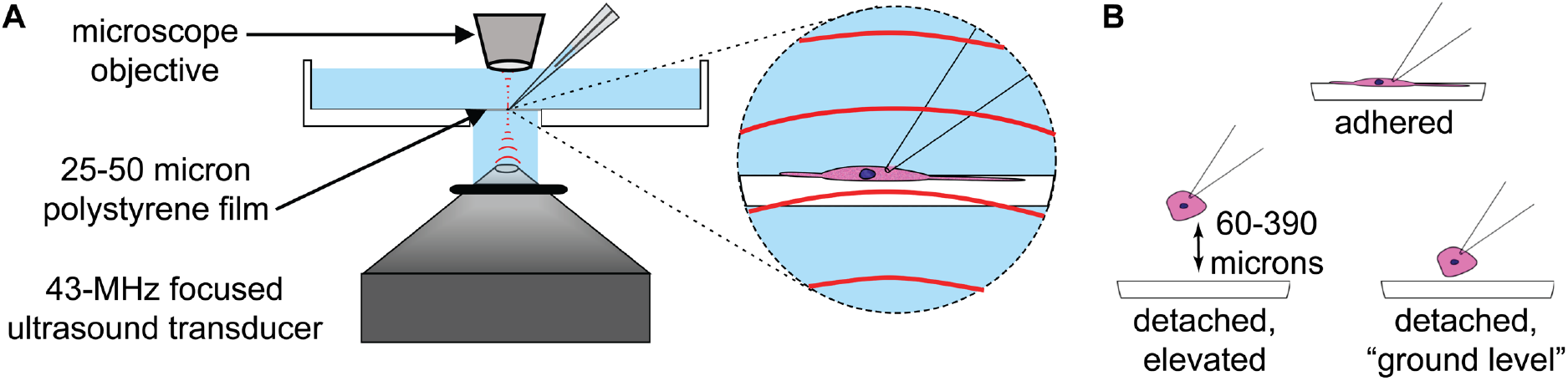
Experimental Set-Up. **A**. Diagram of experimental set-up. Focused ultrasound (US) at 43 MHz is propagated from below to cells adhering to a thin (25- or 50-micron) polystyrene film, oriented perpendicular to the direction of US propagation. See Experimental Procedures for details. **B.** Cartoon illustrating the three different whole-cell recording configurations used in our experiments. Cells were either adhered to the polystyrene film (*top*), or detached and elevated above it (*bottom left*), or detached but not elevated (“ground level,” *bottom right*). The cartoon is not to scale.

### Electrophysiology

Voltage-clamp recordings were performed using an Axopatch 200B amplifier (Molecular Devices, Sunnyvale, CA, USA) with a Digidata 1440 A digitizer (Molecular Devices, Sunnyvale, CA, USA) and pClamp 10.4 software (Molecular Devices, Sunnyvale, CA, USA). Currents were sampled at 100 kHz and filtered at 5 kHz for Piezo1 channels or 10 kHz for Na_V_1.2 channels. Patch-clamp pipettes were pulled from thin-walled glass using a Sutter Instruments P-87 puller (Sutter Instruments, Novata, CA, USA) and had resistances between 2 and 7 MOhm when filled with the internal solution. The internal and external solutions were, respectively, in mM: 125 CsCl, 1 MgCl_2_, 1 CaCl_2_, 4 Na2ATP, 0.4 Na2GTP, 5 EGTA, pH 7.3 (CsOH) and 127 NaCl, 3 KCl, 1 MgCl_2_, 10 HEPES, 10 glucose, pH 7.3 (NaOH) for the experiments on Piezo1 channels; and 140 KCl, 2 MgCl_2_, 1 CaCl_2_, 11 EGTA, 10 HEPES, pH 7.2 (NMDG) and 140 NaCl, 3 KCl, 2 MgCl_2_, 1 CaCl_2_, 10 HEPES, pH 7.2 (NMDG) for the experiments on Na_V_1.2 channels. Series resistance compensation was not used due to the small size of the currents in these experiments. For experiments on Na_V_1.2 channels, data were rejected if the magnitude of the maximum series resistance error (*ΔV*_max_), calculated according to

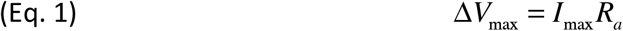

where *I*_max_ is the peak current and *R_a_* is the access resistance, was more than 10 mV. For experiments on Na_V_1.2 channels, pipettes were coated with dental wax to reduce pipette capacitance.

#### Elevated cells

In some experiments, cells were detached and elevated above the bottom of the chamber to examine the effect of height on the response to US. For these experiments, the height of the cell was determined by reading the display on the controller for the micromanipulator (Sutter Instruments MPC-325 (Sutter Instruments, Novata, CA, USA)), or (in experiments using an Eppendorf Patchman micromanipulator (Eppendorf, Hamburg, Germany) lacking this feature) by counting the number of rotations of the focus adjustment knob required to bring the cell into focus starting from the focal plane at the bottom of the chamber, and then converting this value into height using the calibration factor provided by the manufacturer. In contrast to cells at the bottom of the chamber, the gigaOhm seals of elevated cells were sometimes damaged by US stimulation, as evidenced by a sudden and irreversible drop of the apparent membrane resistance to <200 MΩ. This effect, which is most likely a result of acoustic streaming, became more frequent with increased height above the bottom of the chamber. The height at which the frequency of this effect made experiments very challenging was ~400 microns, so experiments were limited to the height range 0-400 microns.

#### Ultrasound

US stimuli were produced by a custom-made focused 43-MHz transducer with a quartz lens. The transducer was calibrated using the method described previously(Prieto, et al. 2013). The focal region of the US beam produced by this transducer is approximately cylindrical with a 90-micron diameter and 500-micron height. Outside of the focal region, the sound beam diverges rapidly, such that the sound reflected from the microscope objective above the cells does not interfere significantly with the sound in the focal region; we therefore do not expect standing waves to be significant in our experimental set-up. This idea was confirmed by finite-element simulations of the acoustic pressure field. The transducer was excited by 43-MHz sine-wave pulses from a Hewlett-Packard 8116A function generator (Hewlett-Packard, Palo Alto, CA, USA), amplified by an ENI 403LA 37-dB amplifier (ENI, Inc., Rochester, NY, USA). The function generator was controlled by an Agilent 33220A function generator (Agilent Technologies, Palo Alto, CA, USA), which was triggered by 5-V pulses from the digitizer. The focal region was localized along the z-axis to the surface of the polystyrene film using a pulse-echo method as described previously (Prieto, et al. 2013). The focal region was localized in the x-y plane by observing the acoustic streaming in response to US using Molecular Probes 1-micron fluorescent beads (Thermo Fisher Scientific, Waltham, MA, USA). A small volume (1-10 microliters) of bead suspension, diluted 1:100 in the external solution, was added to the chamber. In response to US, a circulatory acoustic streaming flow occurs, such that the beads appear to converge on the focal zone. Using the controls intended for centering the condenser, the focal zone was positioned at the center of the microscope’s field of view. The solution in the chamber was then replaced with fresh solution and the cell chosen for recording was placed in the center of the focal region by adjusting the microscope stage. All ultrasound stimuli were continuous wave (100% duty cycle), and intensities are reported as the spatial peak, pulse average intensity at the focus of the ultrasound beam.

### Data analysis

Data were analyzed and curve fitting was performed using Igor Pro (Wavemetrics, Lake Oswego, OR, USA). Ion channel activation and inactivation rates were determined from single exponential fits to the time course of the current rise and decay. All current records analyzed, with the exception of one of the experiments on Piezo1 channels, are the average of at least three trials.

### Experimental design and statistical analysis

Two tailed t-tests were used to establish statistical significance. Significance was defined as P < 0.05. All mean results are reported and displayed as mean ± standard deviation (SD). For the analysis of Piezo1 voltage dependence, one recording with an anomalous current-voltage relationship was excluded based on a Dixon’s Q-test (Barnett and Lewis 1994).

### Finite-elementsimulations

Finite-element simulations of the acoustic pressure, acoustic streaming, and acoustic heating in the experimental chamber during our experiments were performed using COMSOL (COMSOL Inc., Palo Alto, CA, USA). The simulation domain had radially symmetrical geometry, and was 6 mm in the axial direction by between 1 and 5 mm in the axial direction (depending on the property simulated), with a 940-micron diameter by 100-micron height arc on the lower axial boundary of the simulation domain representing the quartz lens of the transducer. The distance from the transducer lens to the bottom surface of the polystyrene was 4.2 mm, and the distance from the polystyrene to the upper boundary of the simulation was 2 mm, corresponding to the focal distance of the microscope objective. For each simulation, an appropriate mesh was determined by iteratively refining the mesh, starting from the default “Very Fine” mesh setting, until changes in the mesh had no effect on the result of the simulation. The default linear or non-linear solver was used for each simulation, except for the acoustic streaming simulation where a segregated Incomplete lower-upper factorization (incomplete LU) solver was used. Due to the radially symmetrical geometry, a symmetrical boundary condition was applied at the inner radial boundary in all simulations. All other boundary conditions are detailed in the descriptions of the individual simulations. Material properties for water and polystyrene used in the simulations, along with sources for these values, are listed in Table 1. The attenuation coefficient for polystyrene was determined by measuring the echoes from the top and bottom surfaces of a 1.6 mm thick sample of the polystyrene used as the cell culture substrate and simulating the sound propagation and echo in COMSOL. Briefly, the amplitude of the reflected pulse from the bottom surface of the sample (the surface closest to the transducer) was used to determine the reflection coefficient between water and polystyrene. The amplitude of the reflected pulse from the top surface was compared with that expected based on this reflection coefficient in the absence of attenuation to determine the attenuation coefficient. We measured the attenuation coefficient at 15, 20, 25, and 30 MHz and extrapolated to 43 MHz assuming linear attenuation with frequency.

**Table 1.**
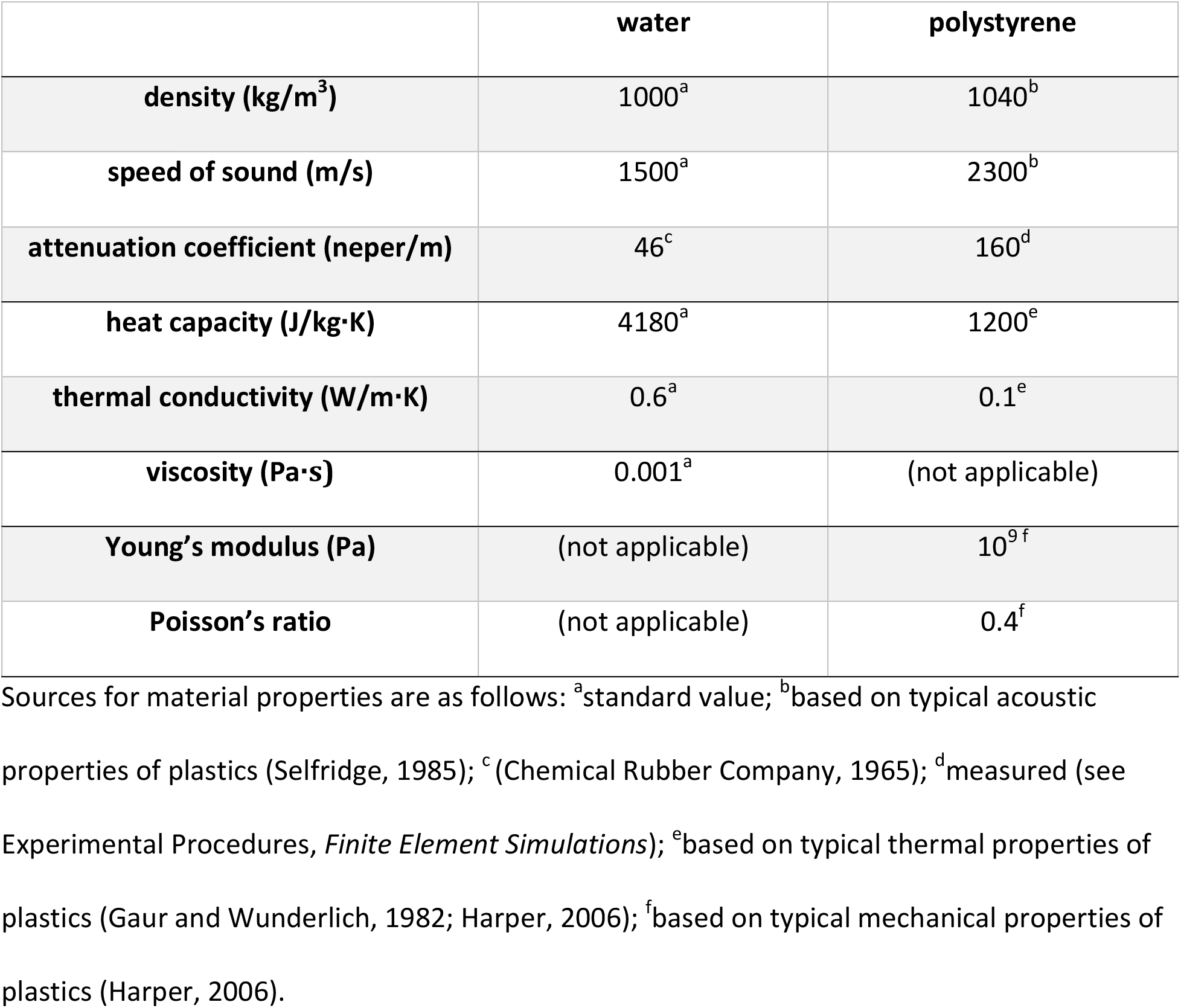
Values of Material Properties Used in Finite-Element Simulations.

|  | **water** | **polystyrene** |
| --- | --- | --- |
| **density (kg/m<sup>3</sup>)** | 1000<sup>a</sup> | 1040<sup>b</sup> |
| **speed of sound (m/s)** | 1500<sup>a</sup> | 2300<sup>b</sup> |
| **attenuation coefficient (neper/m)** | 46<sup>c</sup> | 160<sup>d</sup> |
| **heat capacity (J/kg·K)** | 4180<sup>a</sup> | 1200<sup>e</sup> |
| **thermal conductivity (W/m·K)** | 0.6<sup>a</sup> | 0.1<sup>e</sup> |
| **viscosity (Pa·s)** | 0.001<sup>a</sup> | (not applicable) |
| **Young's modulus (Pa)** | (not applicable) | 10<sup>9f</sup> |
| **Poisson's ratio** | (not applicable) | 0.4<sup>f</sup> |

Sources for material properties are as follows: <sup>a</sup>standard value; <sup>b</sup>based on typical acoustic
properties of plastics (Selfridge, 1985); <sup>c</sup>(Chemical Rubber Company, 1965); <sup>d</sup>measured (see
Experimental Procedures, *Finite Element Simulations*); <sup>e</sup>based on typical thermal properties of
plastics (Gaur and Wunderlich, 1982; Harper, 2006); <sup>f</sup>based on typical mechanical properties of
plastics (Harper, 2006).

#### Simulation of the acoustic pressure field

The steady acoustic pressure field was determined by solving the acoustic wave equation (Pierce 1994)

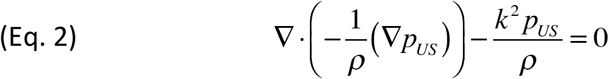

where ρ is the resting density, *p_US_* is the complex acoustic pressure, and *k* is the complex wavenumber defined as

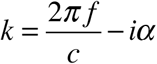

with frequency *f*, speed of sound *c*, and attenuation coefficient *a*. Equation 2 was solved with a 150-micron perfectly matched layer on the outer radial boundary and hard reflecting boundary conditions on all remaining boundaries. An oscillating pressure of 50 kPa or 67 kPa was applied at the boundary corresponding to the transducer lens. The pressure amplitude on the lens was chosen so that the simulated pressure in the focal region, in a simulation performed with the polystyrene film replaced by water, matched the pressures used in the experiments, as determined from the transducer calibration. The radial domain size was 1 mm.

#### Simulation of acoustic streaming

Acoustic streaming at steady-state was simulated by solving the Navier-Stokes Equations for an incompressible fluid with a volume force term due to acoustic radiation force (Nyborg 2008):

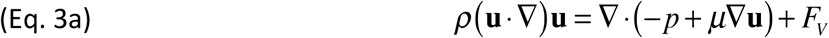

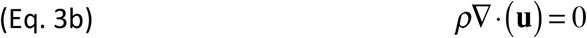

where **u** is the acoustic streaming velocity, *p* is the hydrostatic pressure, *μ* is viscosity, and *F_V_* is the radiation force per unit volume:

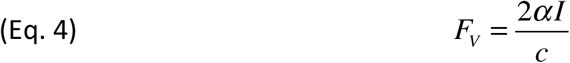

with the US intensity *I* defined as:

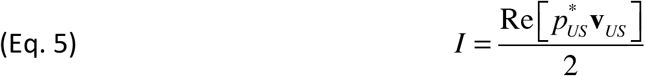

where *p^*^_US_* is the complex conjugate of the complex acoustic pressure and ***v**_US_* is the complex particle velocity, obtained by

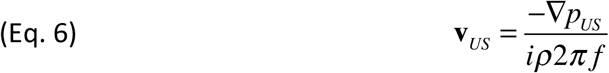

No slip (zero-velocity) boundary conditions were applied at the upper and lower axial boundaries, at the outer radial boundary and at the upper and lower surfaces of the polystyrene film. We also included the effect of radiation force on the polystyrene. The polystyrene buckles slightly under the influence of radiation force, which determines the position of the no-slip boundary. The polystyrene was modeled as an elastic material with geometric nonlinearity included due to the thinness and softness of the material. The displacement was determined by solving:

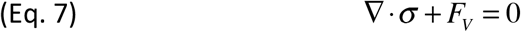

where *σ* is the stress tensor and *F_V_* is the radiation force. The components of the stress tensor are a function of the strain (gradient of the displacement) and of Young’s modulus (E) and Poisson’s ratio (*v*) (Landau and Lifshitz 1986).

The radial domain size for the simulated displacement was 5 mm and the displacement was fixed at zero at the outer radial boundary. The displacement of the polystyrene film was fully coupled to the movement of the fluid using the Fluid-Structure Interaction multiphysics module of COMSOL. Since boundary-layer effects can have very significant impact on fluid dynamics simulations, we performed simulations in which the distance to the radial boundary was changed between 1 and 5 mm. These simulations indicated that the presence of this arbitrary (non-physical) boundary does not affect the simulated streaming within the focal region. We also performed simulations in which the distance to upper axial boundary (a physically meaningful boundary corresponding to the surface of the microscope objective) was changed, to determine whether the position of this boundary (which was changed during experiments on elevated cells due to adjustment of the microscope focus) affects acoustic streaming. These simulations indicated that the position of the upper axial boundary does not affect the streaming in the focal region.

#### Simulation of acoustic heating

Acoustic heating was simulated by solving the heat transfer equation (Hand 1998):

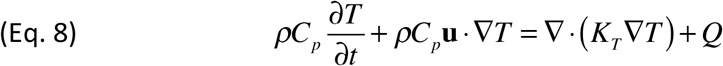

where *C_p_* is the specific heat capacity (at constant pressure), *T* is temperature, **u** is the streaming velocity, *K_T_* is the thermal conductivity, and the heat source *Q* is determined by the US intensity and attenuation coefficient according to *Q* = 2*Iα*. A thermally insulating boundary condition was applied on all boundaries except the inner radial boundary. The radial domain size was 1 mm.

> *Note on heating measurements*. We anticipated that absorption of acoustic energy by polystyrene could cause significant heating in our experiments, which we minimized by using thin (25-50 micron) polystyrene films as the cell-culture substrate rather than standard polystyrene cell-culture dishes. Despite these precautions, a small but significant temperature change occurs during our experiments, although the precise value of the temperature change is difficult to quantify because reliable thermal measurements in the presence of highly focused ultrasound fields are not technically feasible. The peak temperature rise in these experiments is expected to be localized to an area approximately equal in diameter to the focal volume (90 microns), which determines the spatial resolution that would be required for an accurate measurement. Solid-state temperature measurement devices such as thermocouples are available on this scale, but these will reflect and distort the sound field and will therefore not provide an accurate measurement. We attempted to measure temperature changes based on changes in the holding current for an open patch-clamp pipette tip (Shapiro, et al. 2012, Yao, et al. 2009) since this method has the appropriate spatial resolution, but in preliminary attempts we discovered that US would sometimes cause an apparent decrease in the temperature measured using this method, indicating that US can affect open-tip holding current through mechanisms other than heating. Instead, we estimated the heating using simulations, combined with measurement of the US attenuation in polystyrene, as described above.

### Video imaging

Supplementary Movie 1 was acquired using a Hamamatsu Orca Flash 4 microscope camera (Hamamatsu Photonics, Sunnyvale, CA, USA) with Micromanager software (Edelstein, et al. 2010) (University of California, San Francisco, CA, USA) at 100 frames per second with a 10-ms exposure time.

### Cell culture and transfection

Rat Na_V_1.2 channels were stably expressed in Chinese Hamster Ovary (CHO) cells. The CHO/Na_V_1.2 cell line was provided by William Catterall (University of Washington). Mouse Piezo1 channels were transiently expressed in CHO or Human Embryonic Kidney (HEK) cells by Lipofectamine (Thermo Fisher Scientific, Waltham, MA, USA) transfection according to the manufacturer’s protocol. Piezo1 in a vector containing an internal ribosome entry sites (IRES) and enhanced green fluorescent protein (eGFP) was provided by Ardem Patapoutian (Scripps Research Institute, La Jolla, CA). Successfully transfected cells were identified by eGFP fluorescence and were recorded from 2-3 days after transfection. Cultured cells were maintained in an incubator at 37 C with 5% CO_2_. The culture media was Dubelco’s Modified Eagle Medium (DMEM)/F12 (1:1, high glucose, with sodium bicarbonate and L-glutamine) with 10% Fetal Bovine Serum (FBS) and 100 μg/mL penicillin/streptomycin for CHO cells and DMEM (high glucose, with sodium pyruvate and L-glutamine) with 10% FBS and 100 μg/mL penicillin/streptomycin for HEK cells. For transiently transfected cells, antibiotics were omitted and FBS was reduced to 5%. For the stably transfected Na_V_1.2 cell line, 200 μg/mL G418 was included to maintain expression. For recording, cells were trypsinized and dissociated and plated on ~2 cm square pieces of polystyrene film (25 or 50 microns thick) which had previously been treated with atmospheric plasma for 20 minutes in a Harrick plasma cleaner (Harrick Plasma, Ithaca, NY, USA) at the high radio frequency setting to create a positively charged surface for cell adhesion. Polystyrene films with cultured cells were placed in 6-well plates, covered with the standard media, and maintained in the incubator until used (1-3 days after plating). For experiments where the goal was to compare the effects of US on adhered versus detached cells, cells that had been plated at least 24 hours previously and appeared flat and elongated were considered fully adhered. Fully adhered cells could be detached from the polystyrene substrate without damaging the seal by a slow, very gentle back-and-forth movement of the patch-pipette. For experiments on detached cells where it was not essential that the cells be fully adhered at the start of the experiment, we used the following procedure to obtain loosely adhered that cells that could be easily detached without damaging the seal: first, cells were trypsinized and dissociated and a small volume (25-100 microliters) of cell suspension was added to the experimental chamber; the chamber was then placed in the incubator, inside a larger dish with a damp KimWipe to prevent dehydration, for 1-4 hours before recording. All chemicals and cell culture reagents were obtained from Sigma-Aldrich (Saint Louis, MO, USA) or Thermo Fisher (Waltham, MA, USA), unless otherwise indicated.

## RESULTS

We measured the effect of ultrasound on heterologously expressed ion channels using a focused 43-MHz transducer that has previously been shown to potentiate action potential firing in the salamander retina (Menz, et al. 2013), using the experimental set-up illustrated in Figure 1A. US is transmitted from below the experimental chamber through a thin (25 or 50 micron) polystyrene film. We measured the effects of US under three different whole-cell recording conditions (Figure 1B): first, with the cells expressing the channels adhered to the polystyrene film at the bottom of the recording chamber (top); second, with the cells detached from the polystrene substrate and elevated above its surface by between 60 and 390 microns (*bottom left*); and finally with the cells detached from the substrate but not elevated above its surface (“ground level,” *bottom right*). We chose to examine these different conditions because US can affect biological systems through several different modalities and, as examined in more detail below, the strength of these modalities, absolutely and relative to one another, can vary widely depending on the location within the experimental chamber. Specifically, in our experimental set-up the spatial distributions of acoustic heating and acoustic streaming have opposite dependences on height above the surface of the polystyrene film: heating is much more prominent for cells adhered to the film or very close to it than for cells elevated above it, while acoustic streaming is absent at the surface of the film but becomes increases with height. The strength of a third potential modality, acoustic radiation force, decreases with height but varies much less dramatically than heating or streaming over the range examined in these experiments (0-390 microns). Thus, examining the effects of US as a function of height above the bottom of the chamber can help distinguish which modalities are responsible for US effects on ion channels.

As seen in Figure 2, US can activate Piezo1 channels when the cell expressing the channels is elevated above the bottom of the chamber, but not when the cell is at “ground level,” and the Piezo1 current in response to US increases with height. Figure 2A shows an example of the current at -80 mV in response to US at 50 W/cm^2^ in a Piezo1-transfected HEK cell elevated above the bottom of the chamber by 330 microns (*black current trace*) and at “ground level” (0 microns, *gray current trace*), where there is no apparent response to ultrasound. This phenomenon was reproducible, with a detectable current response to US at 50 or 90 W/cm^2^, at a height of 60 microns or higher, in 26 out of 32 cells tested, but never at ground level, whether or not the cell was the adhered to the bottom of the chamber. The effect of height on the US-activated current was reversible (Figure 2B-C), indicating that the sensitivity to US in elevated cells is not due to irreversible damage to the cell rendering Piezo channels more responsive to US (as might occur, for example, if US were to disrupt the cytoskeleton, reducing resting membrane tension and thereby making a previously inactivated population of Piezo channels available for activation by US). The dependence of the peak US-activated current on height is summarized in Figure 2D for twelve different cells for which recordings at multiple heights were obtained. The effect of height on the current amplitude is summarized for all cells in Figure 2E, which shows the distribution of current amplitudes for cells below (*triangles*) or above (*circles*) 50 microns. The US-activated current inactivates with a time constant on the order of that previously reported for heterologously expressed Piezo1 channels (~13 ms (Coste, et al. 2010, Coste, et al. 2012)), although there was significant variability in time constant among cells tested (Figure 2F). Inactivation rates and peak current values are weakly voltage-dependent, as is characteristic of Piezo channels (Figure 2G-I).

**Figure 2.**
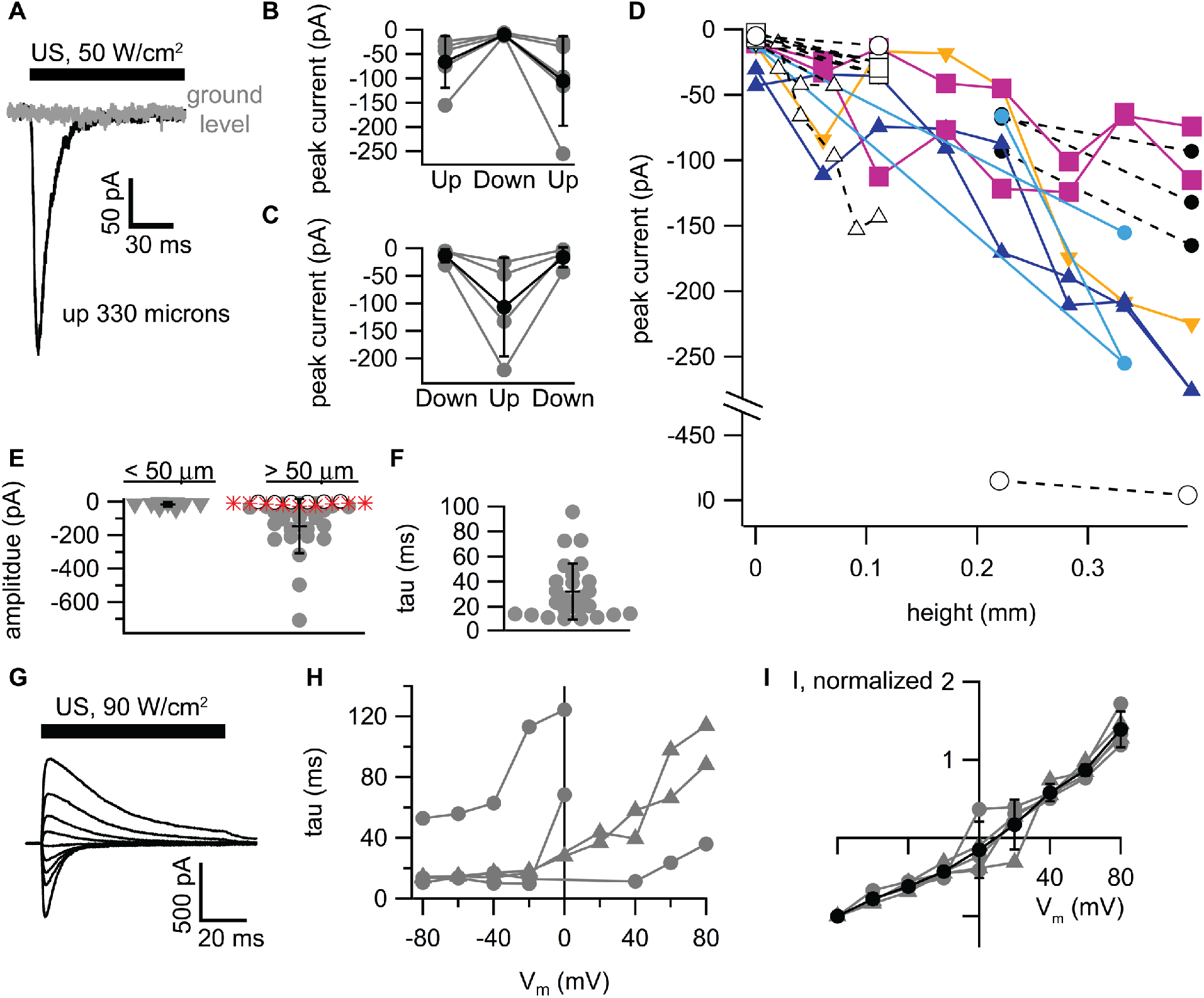
Activation of Piezo1 channels by ultrasound (US). **A.** Example currents at -80 mV in response to a 200-ms, 50 W/cm^2^ US application in a Human Embryonic Kidney (HEK) cell expressing Piezo1 channels, recorded while the cell is at the bottom of the recording chamber (*gray current trace*, “ground level”) and after detaching the cell from film and elevating it by 330 microns (*black current trace*). **B-C.** Reversibility of the effect of height on US-activated currents. Cells were sequentially exposed to US at 50 or 90 W/cm^2^ at an elevated height (70-390 microns, “Up”), then at the bottom of the chamber (“Down”), then back to the original height (B); or at the bottom of the chamber, then at an elevated height, then back to the bottom of the chamber (*C*). Individual cells are shown as gray circles and mean ± standard deviation (SD) values as black circles. N=5 in *B* and N=4 in *C*. **D.** Peak US-activated current as a function of height for N=12 cells. Each group of connected symbols represents one cell tested at various heights. US was either at 50 W/cm^2^ (*small symbols*) or 90 W/cm^2^ (*large symbols*). The different symbol shapes, colors, and line styles are intended to help distinguish individual cells and have no additional symbolic meaning. **E.** Distribution of current amplitudes at -80 mV for cells examined at heights > 50 microns (*circles*, N = 31) or < 50 microns (*triangles*, N = 9, same cells as at > 50 microns; data at lower heights not available in all cells). Open circles indicate Piezo1-transfected cells with no detectable Piezo current (N = 6), and red stars indicate control cells transfected with enhanced green fluorescent protein (eGFP) only, all of which showed no detectable US-activated current (N = 9). For this analysis, data at the highest height available in each of the two categories was used for cells tested at multiple heights. One cell was excluded from this analysis because the height was not quantified. The mean and standard deviation for cells with detectable Piezo1 current are indicated (-16 ± 14 pA, < 50 microns (N = 9); -145 ± 163 pA, > 50 microns (N = 25)) The difference in current amplitude between groups was statistically significant (for all cells: unpaired t-test, unequal variance, P = 0.0006, N = 25; for cells in both groups: paired t-test, P = 0.004, N = 9). **F.** Distribution of inactivation time constants (*tau*) for US-activated currents at -80 mV for N = 26 cells. The mean and standard deviation are indicated (33.1 ± 23.5 ms) **G.** Example currents at a series of voltages from -80 to +80 mV (in 20 mV steps) in a Chinese Hamster Ovary (CHO) cell expressing Piezo1 channels in response to a 200-ms, 90=W/cm^2^ US application. The cell was 170 microns above the bottom of the chamber. **H.** Inactivation time constant for Piezo1 currents expressed in CHO cells (*gray triangles*) or HEK cells (*gray circles*) in response to US at 90 W/cm^2^ as a function of membrane voltage (V_m_) for N=6 different cells. I. Peak US-activated current as a function of membrane voltage (V_m_) for the cells in H, normalized to the current at -80 mV, along with the mean ± SD values (*black circles*).

We considered whether the US-activated current might be due to endogenous mechanosensitive channels. HEK cells express an endogenous mechanosensitive current that can be eliminated by knocking out Piezo1, but this current exhibits very slow inactivation (time constant ≫100 ms) (Dubin, et al. 2017). To confirm that the currents in response to US are due to heterologously expressed Piezo1 channels, we compared currents in Piezo1-transfected cells with those in control cells of the same batch transfected with eGFP in the same expression vector. In nine out nine cells transfected with eGFP alone, there was no apparent US-activated current at 50 or 90 W/cm^2^ at a height of 220 or 390 microns (Figure 2E, *red stars*) compared with seven out of seven cells concurrently transfected with Piezo showing a significant (30 – 500 pA) response to US under the same conditions. Based on these results, and on the relatively fast inactivation time constants of the currents (Figure 2F) compared to those reported for endogenous channels, we conclude that the US-activated current is primarily due to heterologously expressed Piezo1 channels, although we cannot entirely rule out some small contribution from endogenous channels.

A relatively small change in position has a profound effect on the response of Piezo1 channels to US. How can a change in position as small as 100 microns render previously unresponsive channels responsive to US? This distance is equivalent to only about three wavelengths at 43 MHz, and is very small relative to the US attenuation coefficient in water (46 neper/m) (www.ondacorp.com/images/Liquids.pdf). To help explain the change in the sensitivity of Piezo1 channels to US in elevated cells, we performed finite-element simulations of three different modalities by which US may affect ion channel activity: acoustic radiation force, acoustic streaming, and acoustic heating (Figures 3 and 4). Radiation force and acoustic streaming are hypothesized to activate Piezo channels by causing cell membrane stress, while heating typically accelerates ion channel kinetics and can therefore activate ion channels if the temperature rise is sufficiently large or the channel kinetics are especially sensitive to heat. The effects of US on all relevant aspects of the experimental set-up, including the solution in the experimental chamber, the polystyrene film at the bottom of the chamber, and the water coupling the chamber to the transducer, were simulated. Although there is some uncertainty in the absolute values of the simulated field parameters due to estimates of the material properties input to the simulation (see references in Table 1 for the range of material properties for various forms of polystyrene and related polymers), our conclusions are based primarily on the spatial distributions of the field parameters, which are not affected by small changes in material properties, rather than their absolute values.

**Figure 3.**
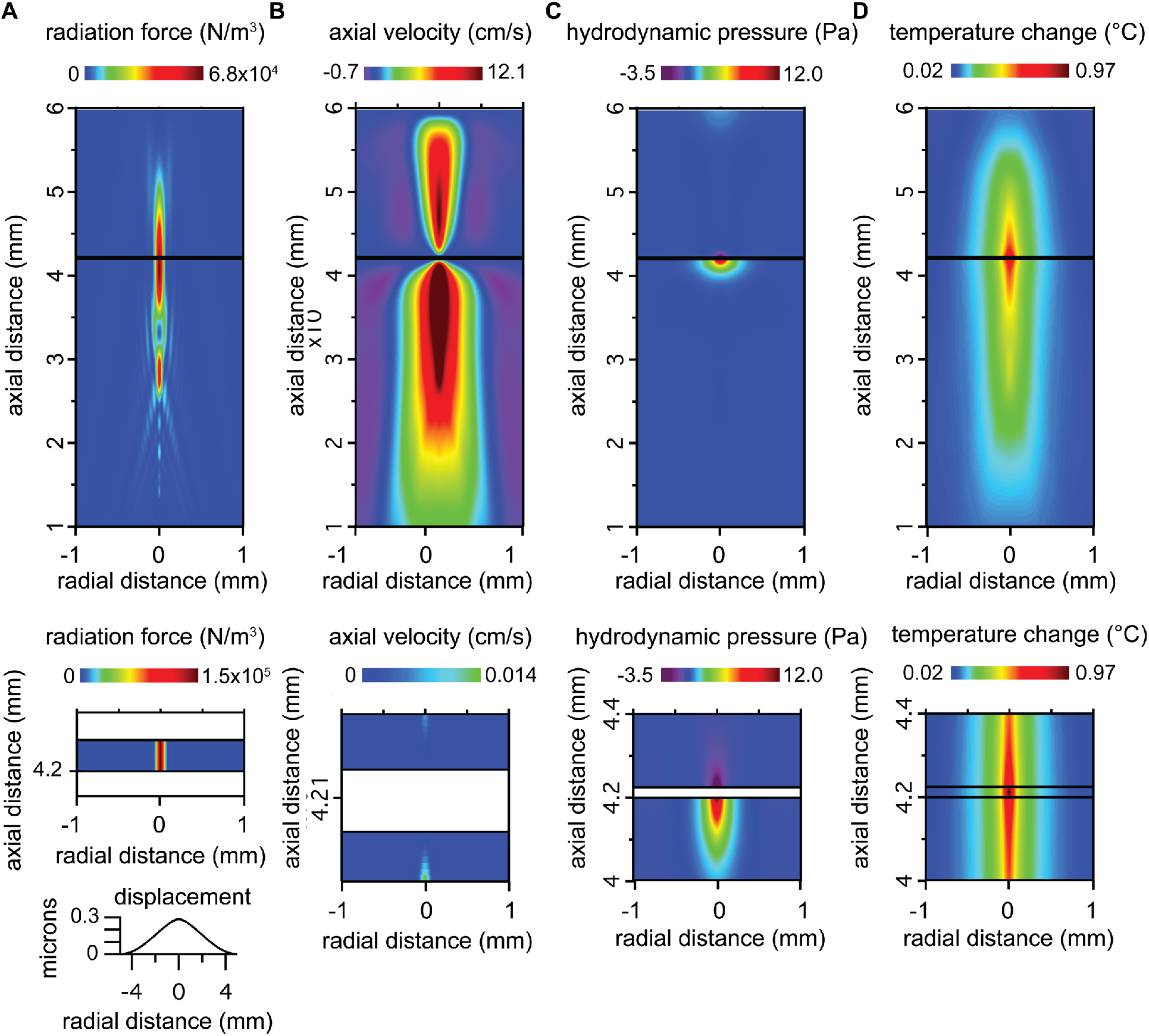
Simulated spatial profiles of acoustic field parameters. **A.** Acoustic radiation force in the liquid media (*top panel*) and in the polystyrene film (*middle panel*). The bottom panel shows the simulated displacement of the surface of the polystyrene film in response to acoustic radiation force. **B.** Axial component of the acoustic streaming velocity. The bottom panel shows the streaming velocity on an expanded scale, highlighting relevant features in the vicinity of the polystyrene film. **C.** Hydrodynamic pressure due to acoustic streaming (shown on an expanded scale in the bottom panel). **D.** Steady-state temperature change (shown on an expanded scale in the bottom panel). Axial distances are relative to the surface of the transducer and radial distances are relative to the center of the ultrasound (US) beam. Results are shown for US at 90 W/cm^2^ and 25-micron polystyrene film. Similar results were obtained for US at 50 W/cm^2^ and 50-micron polystyrene (not shown). Changing the US intensity or the thickness of the polystyrene film changed the amplitude of the acoustic field parameters but did not substantially change their spatial profile.

**Figure 4.**
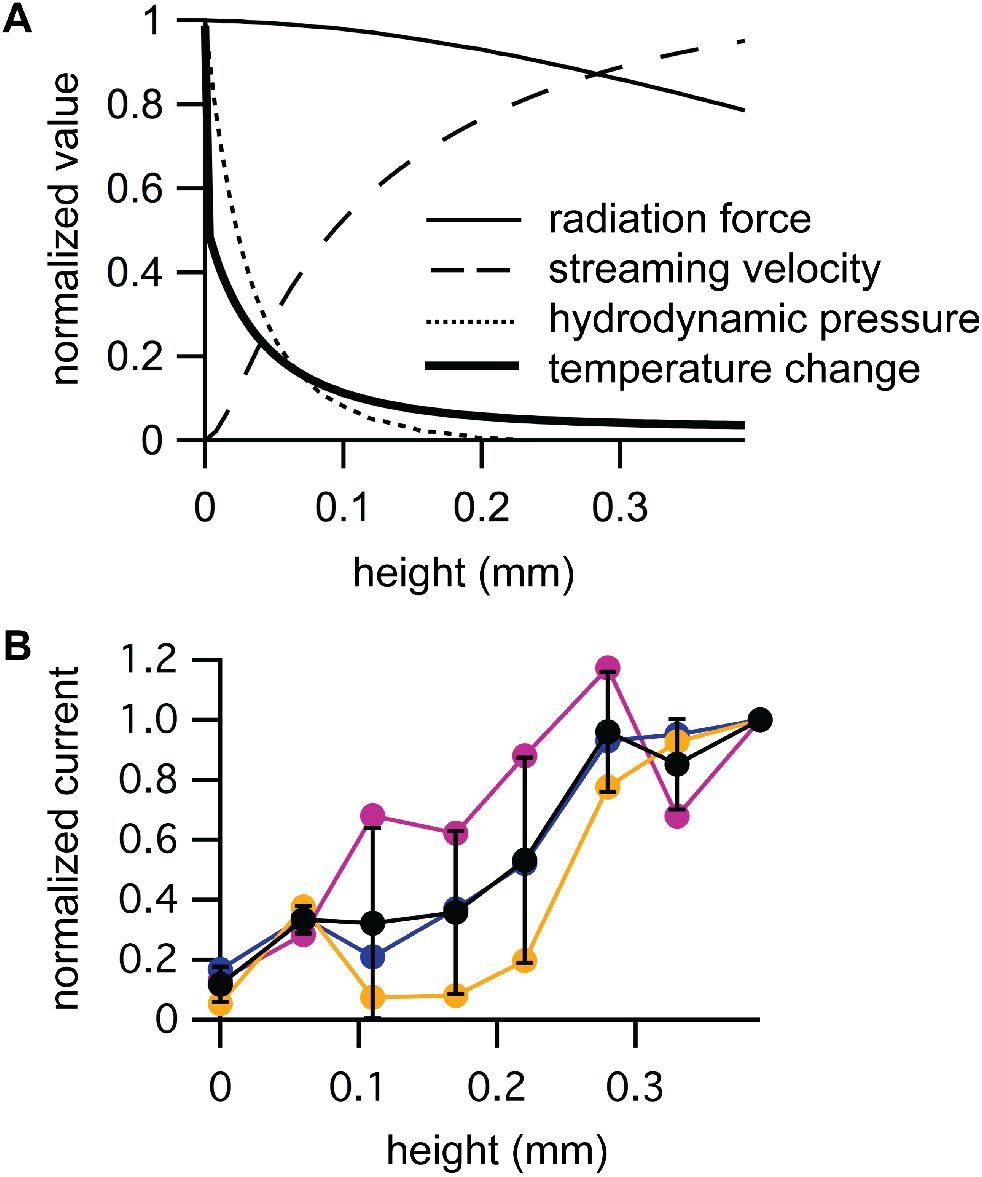
Activation of Piezo1 channels by ultrasound (US) depends on acoustic streaming velocity. **A.** Normalized axial values of the simulated acoustic radiation force (*thin solid line*), acoustic streaming velocity in the axial direction (*dashed line*), hydrodynamic pressure amplitude (*dotted line*), and steady-state temperature change (*thick solid line*) as a function of height above the bottom of the experimental chamber. B. Normalized peak Piezo1 current at - 80 mV in detached cells in response to US as a function of height above the bottom of the experimental chamber, normalized to the peak current at a height of 390 microns for N = 3 cells (*colored circles, corresponding to the colors in Figure 2D*), along with the mean (± standard deviation (SD)) values (*black circles*).

The simulated spatial profile of radiation force in response to ultrasound at 90 W/cm^2^ is shown in Figure 3A for the liquid media (*top panel*) and for the polystyrene film (*middle panel*). Standing waves could create variations in US intensity with spatial period equal to the wavelength, possibly explaining the variation in response to US with height, but the simulation indicates that there is no significant effect of standing waves on the US intensity distribution within the chamber, as there is no apparent periodic variation in radiation force, which is proportional to the US intensity. Based on the simulation, cells at ground level experience a radiation force per unit volume of ~6 N/m^3^, but this is apparently not effective, in and of itself, in activating Piezo1 channels. More importantly, the spatial profile of radiation force is incompatible with direct activation of Piezo1 channels by radiation force, since radiation force decreases with height above the bottom of the chamber (Figure 4A, *thin solid line*), while the response of Piezo1 channels to US increases (Figure 2D). We also simulated the displacement of the polystyrene film in response to radiation force (Figure 3A, *bottom panel*), because a large displacement of the film could affect the spatial distribution of the acoustic field parameters, and a gradient in the displacement could cause stress in cells adhered to the film. The peak displacement is very small (0.3 microns) indicating that displacement of the polystyrene film is not a significant factor in our experiments.

In contrast, the simulated spatial profile of the axial component of the acoustic streaming velocity correlates well with the dependence of the US-activated Piezo1 current on height (Figure 3B). As typically seen with acoustic streaming (Duck 1998), high-velocity streaming in the direction of propagation occurs within the US beam, along with a slower, circulatory return flow outside the beam, as indicated by the regions with negative velocity. (The circulatory flow also has a radial velocity component, much smaller than the axial component (9.4 mm peak radial velocity amplitude), not shown in Figure 3B). Critically, there is a thin stagnant layer, directly above the surface of the polystyrene film, in which no streaming motion occurs (Figure 3B, *bottom panel*). This stagnant layer is the result of the zero-velocity boundary condition at the rigid surface of the polystyrene film. Outside the stagnant layer, the axial streaming velocity increases steeply with height and then levels off over the range probed in the Piezo1 experiments (Figure 4A, *dashed line*). This result explains why Piezo1 channels in cells at the bottom of the chamber are immune to US stimulation, despite the significant radiation force, and why the Piezo1 current in response to US increases with height. We conclude that forces acting on the cell membrane as a result of acoustic streaming are sufficient to activate Piezo1 channels. These forces are caused by relative motion between the cell, which is held in place by the patch-pipette, and the surrounding fluid. A cell suspended in a moving fluid will experience a drag force made up of normal and shear components (King 2002), resulting in membrane stress. Consistent with this idea, we observed that the membranes of elevated cells are distorted and apparently displaced in the axial direction during US stimulation (Supplementary Movie 1). Thus, radiation force is critical for the response of Piezo1 channels to US, insofar as it drives acoustic streaming, but the boundary conditions that regulate acoustic streaming are equally critical.

Although the spatial profile of acoustic streaming velocity appears sufficient to explain our results on Piezo1 channels, we considered whether hydrodynamic pressure caused by streaming might also contribute to the effects of US on Piezo1 channels. The simulated spatial distribution of the hydrodynamic pressure (Figure 3C) indicates that there is a significant negative hydrodynamic pressure within the chamber at the surface of the polystyrene film. However, the amplitude of this hydrodynamic pressure decreases with height (Figure 3A, *dotted line*), which is again inconsistent with the dependence of US-activated Piezo1 current on height. We therefore conclude that, relative to the streaming velocity, hydrodynamic pressure is not an important factor in the activation of Piezo1 channels by US.

Finally, the simulated spatial profile of acoustic heating indicates that temperature changes due to US absorption cannot explain the effects of US on Piezo1 channels. The simulated spatial profile of the temperature rise at the end of a 1-s US stimulus at 90 W/cm^2^ (during which the temperature change reaches an approximate steady state) shows that US-induced temperature rise is greatest at center of polystyrene film (Figure 3D), as expected based on the higher ultrasound attenuation coefficient and lower specific heat of polystyrene relative to water. In contrast to the effect of height on the response of Piezo1 channels to US, the temperature rise steeply declines with height (Figure 4A, *thick solid line*). We therefore conclude that heating does not play a significant role in the response of Piezo1 channels to US. This conclusion is consistent with the observation that mechanically activated Piezo current have unexceptional temperature sensitivity, as reflected in their Q10 temperature coefficient of ~2.8 for inactivation (A. Patapoutian and B. Coste, personal communication).

To further support our conclusion that the activation of Piezo1 channels by US correlates with acoustic streaming, we compared three sets of recordings in which we measured US-activated currents from the same cell across a wide range of heights (Figure 4B). The peak Piezo1 current in response to US increases from its minimum value at zero height to a maximum value at 390 microns, the highest level tested, where it appears to approach a plateau. At this level and above, patch-clamp seals were frequently irreversibly damaged by US, presumably due to acoustic streaming, so we were unable to confirm the presence of a plateau in the Piezo current response. Nonetheless, the height dependence of the US response in Piezo-transfected cells over the range tested clearly resembles the axial profile of the acoustic streaming velocity, which rises from zero at the polystyrene surface to a plateau value at above 300 microns, rather than that of the radiation force, which remains relatively constant between zero and 400 microns, or the hydrodynamic pressure, which declines as the velocity increases, or the temperature change, which rapidly declines from its peak value at the zero microns to a small fraction of the peak at 200 microns (Figure 4A). This result confirms that acoustic streaming is critical for activating Piezo channels with US in our experiments.

We also considered whether, and through what modality, US could affect voltage-dependent Na_V_1.2 sodium channels. Since US can potentiate action potential firing and Na_V_ channels are central to action potential firing, and since the kinetics of least some subtypes of Na_V_ channels can be modulated by membrane stretch (Beyder, et al. 2010, Morris and Juranka 2007, Wang, et al. 2009), it seemed possible that these channels might also respond to US. We therefore measured the effects of US on heterologously expressed Na_V_1.2 channels in cells at ground level and in elevated cells. A representative example of the effect of US at 43 MHz and 90 W/cm^2^ on Na_V_1.2 channels in an adhered cell is shown in Figure 5A. The current in response to a step from a holding potential of -100 mV to -10 mV is shown with (*black current trace*) and without (*gray current trace*) US stimulation. There was no significant effect of US on the amplitude of the Na_V_1.2 current in adhered cells (-379 ± 159, control, versus -382 ± 150 pA, US; mean ± SD, N =6, P = 0.87, paired t-test)). However, the macroscopic rates of activation and inactivation both increase in adhered cells in the US condition, from 2420 ± 530 s^-1^ to 3280 ± 820 s^-1^ (activation) and from 600 ± 140 s^-1^ to 740 ± 150 (inactivation) (mean ± SD), N = 6). The effect of US was significant for both activation (paired t-test, P = 0.006) and inactivation (paired t-test, P = 2×10^−5^). These accelerated kinetics are consistent with either a thermal or a radiation force mechanism. Based on our finite-element simulations, the US-induced temperature rise is 0.8 C at the surface of the polystyrene film at the start of voltage step. For a Q10 of 3, which is typical of ion channel kinetics and similar to that reported for Na_V_ currents (Frankenhaeuser and Moore 1963, Hodgkin, et al. 1952), this would produce a 1.09-fold acceleration of channel gating kinetics, slightly lower than the experimental result (1.23 ± 0.06). However, the exact value of the temperature rise in the simulation depends on the specific heat and thermal conductivity of the polystyrene film, for which we lack exact values. The effect of US on Na_V_1.2 kinetics was not dependent on the size of the voltage step used to activate the channels (data not shown), indicating that the effect of US is orthogonal to the effect of voltage, consistent with a temperature effect. However, membrane stretch produced by applying pressure to cell-attached membrane patches also increases the macroscopic rates of activation and inactivation in Na_V_ channels (although the stretch sensitivity of the specific subtype studied here has not to our knowledge been tested). Thus, it is difficult to tell based on these results alone whether the effects of US on Na_V_ channels are due to acoustic heating, or membrane stress caused by radiation force or acoustic streaming, or some combination of these.

**Figure 5.**
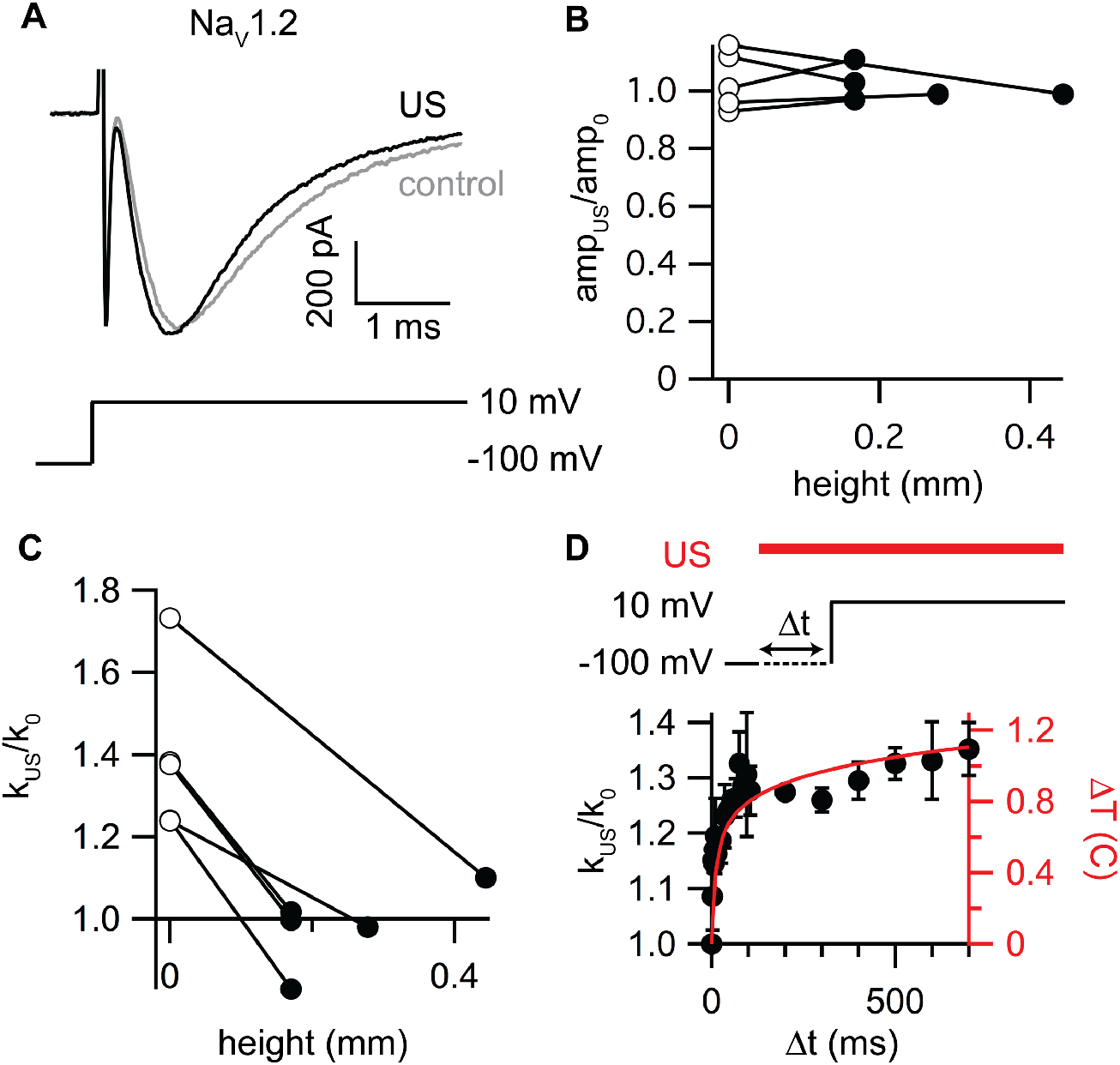
Effects of ultrasound (US) on Nav1.2 Channels. **A.** Example currents for NaV1.2 channels (in adhered cells at ground level) in response to a voltage step from -100 to -10 mV with (*black current trace*) and without (*grey current trace*) stimulation. The 90-W/cm^2^, 1-s US stimulus was applied starting 800 ms before the voltage step. B. Ratio of the Na_V_1.2 current amplitude at -10 mV in the US condition to the amplitude in the control condition (*amp_US_/amp_0_*) for five detached cells at zero height (*open circles*) and at various heights above the bottom of the experimental chamber (*black circles*), for the same stimulus protocol used in A. There was no significant effect of height on the amplitude ratio (paired t-test, P = 0.71). C. Ratio of the Na_V_1.2 inactivation rate at -10 mV in the US condition to the inactivation rate in the control condition (*k_US_/k_0_*) in five detached cells at zero height (*open circles*) and at various heights above the bottom of the experimental chamber (*black circles*), for the same stimulus protocol used in part A. The difference was statistically significant (paired t-test, P = 0.001, N = 5) D. Comparison of the time courses of simulated acoustic heating and effects of US on Na_V_1.2 channel inactivation. The mean (± standard deviation (SD)) ratio of the Na_V_1.2 inactivation rate at -10 mV in the US condition to the inactivation rate in the control condition (*k_US_/k_0_*) is shown as a function of the US exposure time (Δt) prior to the voltage step for N = 4 different cells at each interval, along with the simulated temperature change as a function of the time relative to the start of the US stimulus (*red line*, right axis). The experiments and simulation were done with 50-micron polystyrene.

To determine the physical basis of the effect of US on Na_V_ channel kinetics, we measured the dependence of this effect on height above the bottom of the experimental chamber and, applying the rationale already established for Piezo channels, compared this dependence with the simulated axial profiles of radiation force, acoustic streaming velocity, hydrodynamic pressure, and steady-state temperature change (Figure 4A). As for adhered cells, there was no significant effect of US on the current amplitude in either detached cells at ground level, or in detached and elevated cells (Figure 5B). In contrast to the effect of US on Piezo1 channels, however, the effect of US on the kinetics of Na_V_1.2 channels declines with height (Figure 5C), and the steep decline matches the axial profile of the US-induced temperature change (Figure 4A, *thick solid line*). Thus, unlike for Piezo1 channels, elevating the cell expressing the channels does not reveal a previously suppressed mechanical response to US in Na_V_1.2 channels. This result indicates that the response of Na_V_1.2 channels to US is exclusively or predominantly due to heating. As a final test of this conclusion, we compared the time course of the effect of US on the Na_V_1.2 inactivation rate with the simulated time course of the temperature change at the surface of the polystyrene film. Figure 5D shows the ratio of the NaV1.2 inactivation rate during the US stimulus to the inactivation rate in the control condition, as a function of the time between the start of the US stimulus and the start of the channel-activating voltage step (Δt). The simulated time course of the US-induced temperature change is plotted on the right axis for comparison. As expected, the time courses are very similar. We therefore conclude that the acceleration of channel gating by US is predominantly due to heating, with little, if any, contribution from membrane stress, either due to radiation force directly or to acoustic streaming. Since this heating is dependent on US absorption by the polystyrene film, the response of Na_V_1.2 channels to US in these experiments is of limited relevance in the context of US effects on brain and excitable tissues *in vivo*. The absence of a mechanical effect, despite the hypothesized mechanical basis of US neuromodulation and the sensitivity of Na_V_ channels to membrane stretch in other contexts, is the most important result of our experiments on Na_V_ channels.

## DISCUSSION

### Comparison with previous US/electrophysiology experiments

Here we report stable patch-clamp recordings in the presence of ultrasound. Previously, Tyler and coworkers reported that patch-clamp seals were unstable in the presence of US, and therefore concluded that patch-clamp recording was not a practical technique for investigating the bio-effects of US (Tyler, et al. 2008). The 43-MHz frequency and 90 W/cm^2^ intensity used in our experiments are both much higher than those used by Tyler and co-workers (440 kHz and 2.9 W/cm^2^), suggesting that relatively high-frequency US may be critical for compatibility with patch-clamp recording. In addition, the US beam in our experiments is very tightly focused, which may contribute to our success at combining US stimulation with patch-clamp recording. Finally, another potentially important factor is the absence of significant standing wave effects, which were minimized by the tightly focused US field.

Kubanek and coworkers reported on effects of pulsed US at 10 MHz on ion channel currents recorded using two-electrode voltage-clamp (TEVC) in *Xenopus* oocytes (Kubanek, et al. 2016). They observed a small increase in Na_V_1.5 conductance in response to US but were unable to resolve the peak currents or kinetics due to the limited clamp speed of TEVC. They did not differentiate between possible sources of US effects; however, they did measure a significant temperature rise that could explain the effects they observed and that is consistent with our findings on Na_V_1.2 channels.

### The radiation force hypothesis of US effects on ion channels

The goal of these experiments was to better understand the mechanisms by which US affects the activity of excitable tissues, and to test the hypothesis that US affects ion channel activity by a mechanical effect on cell membranes. While this idea has been proposed by several groups, a correlation between the amount of acoustic radiation force, the amount of membrane stress, or other relevant parameter, and the strength of the response to US has not been demonstrated previously. In fact, US neuromodulation of mouse motor cortex *in vivo* appears to be more effective at lower frequencies (King, et al. 2013, Tufail, et al. 2010, Ye, et al. 2016). This is the opposite of the expected frequency dependence for a radiation force mechanism, since radiation force is proportional to the attenuation coefficient, *a* (Eq. 4), which increases with acoustic frequency. However, it is not yet clear to what extent confounds such as differences in beam profile and absorption by the skull distort the frequency response in *in vivo* experiments. US neuromodulation of the salamander retina *in vitro* is more effective at higher frequency, as expected for a radiation force-based mechanism (Menz, et al. 2016). In heart tissue, Dalecki et al. demonstrated that aortic pressure in the frog heart is modulated by US, and by varying both frequency and beam width, determined that the effect of US was proportional to radiation force, providing compelling evidence for a radiation force mechanism for US modulation of electrical activity (Dalecki, et al. 1997).

Our results on Piezo1 channels are consistent with the general idea that radiation force is involved in the effect of US on electrical activity, insofar as acoustic streaming represents the response of liquid media to radiation force, and can therefor occur at other frequencies besides the 43 MHz frequency tested here. We also demonstrate that US can activate mechanosensitive ion channels, a key component of the radiation force hypothesis. However, critical issues must be resolved to apply these results to *in vivo* neuromodulation experiments. First, the 43-MHz frequency used in our experiments is much higher than the typical frequency range for *in vivo* experiments (~0.2-3 MHz). The driving force for acoustic streaming increases is acoustic radiation force, which increases with frequency due to the frequency dependence of the attenuation coefficient. The relationship between streaming velocity and attenuation is not straightforward because of the importance of geometry and nonlinearity in fluid dynamics, but geometry aside, streaming velocity is expected to increase with frequency, the opposite of the frequency dependence of *in vivo* US neuromodulation responses. However, as already mentioned, there is an apparent discrepancy between the frequency dependence of *in vivo* and *in vitro* US neuromodulation experiments. We cannot rule out the possibility that effects based on cavitation that are not present in our experiments are important at lower frequencies. In addition, methods to evaluate acoustic streaming effects in brain tissue will need to be developed. Relatively simple viscoelastic models of US-tissue interactions do not account for acoustic streaming. US-induced tissue displacement, produced through the same radiation force mechanism that drives acoustic streaming, is the closest analogue to acoustic streaming in these models (Sarvazyan, et al. 2010). Dual-phase models of tissue as a fluid-saturated porous solid will be more appropriate for modeling effects of acoustic streaming.

### Comparison with in vitro neurostimulation experiments at 43 MHz

There is a very interesting parallel between our experiments and those of Menz and Baccus, who used the same focused 43-MHz transducer to stimulate neural activity in the salamander retina *in vitro*, while recording the activity of retinal ganglion cells using a multi-electrode array (Menz, et al. 2013). They found that retinal ganglion cells do not respond directly to US, but show increased activity due to US stimulation of cells in more superficial layers. A key feature of their experiments in the context of the present results is that the ganglion cell layer is directly above the rigid surface of the multi-electrode array, which therefore enforces a no-slip boundary condition at its surface, just as the polystyrene substrate does in our experiments. Thus ganglion cells may be unresponsive to US in these experiments not for reasons related to their biophysical properties, but instead because they are located in a stagnant layer adjacent to the surface of the array. Consistent with this idea, they found that the membranes of ganglion cells undergo much less displacement than those of neurons in more superficial layers in response to US stimulation (Menz, et al. 2013). Together with the experiments reported here, and the observation of neuromodulatory effects in rat hippocampal brain slices *in vitro* (Prieto, et al. 2016), these results motivate further investigation of the neuromodulatory effects of higher frequency US and of the role of acoustic streaming in these effects. In fact, apparent membrane strain and changes in membrane permeability have been observed in single cells in response to stimulation with highly focused ultrasound beams at 200 MHz (Hwang, et al. 2016, Hwang, et al. 2012, Hwang, et al. 2014), suggesting that a mechanism similar to the one we propose here may be occurring at this very high frequency. Since many tissues express mechanosensitive ion channels (Volkers, et al. 2015, Wu, et al. 2016), the scope for useful clinical applications is quite broad if the effects reported here can be reproduced in the clinical setting.

### Absence of a significant mechanical effect on Na_V_1.2 channels

Although we cannot entirely rule out the idea that some small part of the response of Na_V_1.2 channels to US is due to mechanical effects, most of the response is clearly due to heating. Why then don’t Na_V_1.2 channels respond to US when subjected to an acoustic streaming velocity sufficient to activate Piezo channels (Figure 5C), despite their reported mechanosensitivity? A simple explanation would be that more membrane tension is required to affect Na_V_1.2 kinetics than is required to activate Piezo1 channels. This explanation is difficult to rule out. The tension required for half-maximal activation of Piezo1 channels is 2.7 mN/m in cell-attached patches (Lewis and Grandl 2015), but the tension required to modulate Na_V_ kinetics is not known. Instead, quantitative comparison of the effects of tension on Na_V_ and Piezo channels must be based on the pressures applied to membrane patches, which is problematic since membrane tension depends on both the applied pressure and the patch’s radius of curvature (Sokabe, et al. 1991), causing considerable variation in the tension produced by a given pressure step (Moe and Blount 2005). Comparison of pressure values without information on patch curvature is therefore extremely approximate, and the observation that pressures required to activate Piezo channels and modulate NaV channels in on-cell patches are similar (on the order of -10 to -40 mmHg) (Bae, et al. 2013, Beyder, et al. 2010, Coste, et al. 2012, Lewis and Grandl 2015, Morris and Juranka 2007)) is not conclusive.

Another potential explanation for the lack of a mechanically-mediated response to US in Na_V_1.2 channels is that Na_V_ channel mechanosensitivity was demonstrated in cell-attached membrane patches, and the mechanical environment of the ion channels in the membrane patch may be different from that of the remaining channels in the cell membrane (Suchyna, et al. 2009, Ursell, et al. 2011). A third alternative is that Na_V_ channels and Piezo1 channels may respond to different forms of membrane stress through different mechanisms, and the fluid stress from acoustic streaming may act through one of these mechanical pathways but not the other. Broadly speaking, mechanosensitive ion channels are thought to respond to membrane stress through lipid bilayer tension (*e.g*., reconstituted Piezo channels (Syeda, et al. 2016), bacterial mechanosensitive channels (Msc channels) (Sukharev, et al. 1994), and mammalian two-pore-domain potassium-selective channels (K2P) channels (Brohawn, et al. 2014)) or interactions with the membrane-associated cytoskeleton (*e.g*., No Mechanoreceptor Potential C (NOMPC) channels (Zhang, et al. 2015) and members of the epithelial sodium channel (ENac) family (Cueva, et al. 2007). Which of these mechanisms is most important for Na_V_ channels is not known.

### The role of the membrane

Since Piezo channels are thought to respond to membrane stress through lipid bilayer tension (Syeda, et al. 2016), it follows that the response to US could be modulated by the mechanical properties of the membrane and therefore by its lipid composition. This may in part account for the variability in the amplitude of the US-induced Piezo channel current. The simplest way of conceptualizing the effect of lipid composition on the response to membrane stress is in terms of an area elastic constant, K_A_, with a change in the membrane area in response to stress resulting in tension (*γ*) in the plane of the membrane according to *γ* = (ΔA/A)K_A_, where A is the resting area and ΔA is the change in area. Ka and the resulting tension are expected to depend on lipid composition (Helfrich 1973). The interplay between lipid composition and sensitivity to shear stress has been observed in activation of G-protein coupled receptors (Gudi, et al. 1998). Another possibility is that the phase state of the membrane can be affected by US. In fact, absorption of acoustic energy by lipid bilayers has been shown to increase at temperatures near the lipid phase transition temperature (Tata and Dunn 1992), and the phase state of lipid bilayers has been shown to modulate ion channel activity (Seeger, et al. 2010). This adds another layer of complexity to the modulation of the US response by membrane lipid composition. Unfortunately, there is no simple way for us to manipulate the membrane lipid composition, and our experiments do not directly report on the effects of US on membrane properties, but rather indirectly indicate that these changes may be taking place through the observed effects on Piezo channels, so we are unable at this time to test these interesting ideas.

### Limitations of the experimental and computational approach

Our patch-clamp approach allows us to directly measure the effects of US on ion channel currents, but beyond its role as a current measurement device, the patch-clamp pipette has additional physical interactions with the cell and the US field that must be accounted for. First, the cell membrane is tightly attached to the pipette. Clearly, a cell suspended in solution would be carried along with the acoustic streaming flow field, and would therefore experience a completely different set of forces than one anchored to a patch pipette. In this sense, the pipette plays a role in determining the response to US. However, cells in biological tissue are not suspended but rather tightly adhered to the surrounding extracellular matrix. Thus, in our interpretation the pipette acts as a “surrogate extracellular matrix,” anchoring the cell in place and allowing membrane strain to occur in response to acoustic streaming. The pipette also interacts with the US field in ways not included in our simulation. Due to the high acoustic impedance between the glass and the surrounding solution, the tip of the pipette will scatter sound from its surface. However, the diameter of the tip (~1 micron) is small relative to the US wavelength, and the scattering will therefore not significantly affect the US intensity distribution. The pipette may also interact with the acoustic streaming field. Boundary layer effects at the pipette surface may create microstreaming and eddy currents around the pipette. The velocity of these microstreaming effects will be proportional to the macroscopic streaming velocity. Thus the relative importance of macroscopic versus microscopic streaming effects in the response of Piezo channels to US cannot be determined from our results.

## CONCLUSIONS

The experiments reported here provide insight into the molecular and biophysical basis of US-induced bio-effects in the nervous system and other tissues. Our results are consistent with the idea that mechanical forces associated with US can activate mechanosensitive ion channels, but a subtle and key point is that radiation force alone is not sufficient. Instead, we conclude that acoustic streaming produced by radiation force is required to activate Piezo1 channels in our experiments (continuous wave US at 43 MHz, 50-90 W/cm^2^), and this leads us to propose that acoustic streaming or radiation force-induced tissue displacement may be required for the neuromodulatory effects of US seen in various *in vivo* and *in vitro* preparations. For both streaming and tissue displacement, boundary layer effects will suppress the response near the boundary. Thus cultured cells adhering to a rigid surface will not provide an appropriate model system for understanding the effects of US on ion channels. Fortunately, the approach described here can be readily applied to other ion channels, providing a platform for investigating the endogenous response of ion channels to US or developing genetically engineered US-sensitive actuators for “sonogenetics.”

## SUPPLEMENTARY DATA

**Supplementary Movie 1. Cell membrane distortion in response to ultrasound.** A CHO cell at the end of a patch pipette at a height of 170 microns above the bottom of the chamber is exposed to US at 50 W/cm^2^ for 200 ms. The direction of US propagation is normal to the field of view. The image was acquired at 100 frames per second and playback is at 7 frames per second. The length of the video is 500 ms.

## ACKNOWLEDGMENTS

This work was funded by National Institute of Health grant 1R01EB019005 (MM, BKY) and by an American Heart Association Western Affiliates postdoctoral fellowship (MLP) and by the Mathers Foundation. The content is solely the responsibility of the authors and does not necessarily represent the official views of the National Institutes of Health. Funding agencies had no role in the decision to publish the results.

We thank Michael Menz for critically reading the manuscript, and members of the Stephen Baccus, Kim Butts-Pauly, and Miriam Goodman labs for helpful discussions. We thank the Steven Boxer lab for sharing their plasma cleaner and the Goodman lab for the loan of their microscope camera.

